# Sensitive and specific miRNA *in situ* hybridization using partially methylated phosphotriester antisense DNA probes

**DOI:** 10.1101/2022.06.12.495852

**Authors:** Po-Hsiang Wang, Tony Z. Jia, Ching-Wen Chang, Bertrand Chin-Ming Tan, Ya-Hui Chi, Wen-Yih Chen

**Affiliations:** Graduate Institute of Environmental Engineering, National Central University, Taoyuan, Taiwan; Earth-Life Science Institute, Tokyo Institute of Technology, Tokyo, Japan; Blue Marble Space Institute of Technology, Seattle, WA, USA; Department of Chemical Engineering and Materials Engineering, National Central University, Taoyuan, Taiwan; Department of Biomedical Sciences, Chang Gung University, Taoyuan, Taiwan; National Health Research Institutes, Miaoli, Taiwan

**Keywords:** Antisense oligonucleotide probe, neutralized DNA, modified nucleic acid, *in situ* hybridization, microRNA, colorectal cancer

## Abstract

Neutralized DNA (nDNA) is an emerging class of DNA oligonucleotides chemically synthesized with site-specific internucleoside methyl phosphotriester linkages, changing the negatively charged DNA phosphodiester backbone to a neutral methyl phosphotriester backbone. The reduction of inter-strand charge repulsion of nucleotide duplexes results in stronger binding between nDNA and other nucleic acids, and as such, nDNA has been used as a sensitive antisense probe for sequencing nucleotides. From a thermodynamic perspective due to steric effects, a hybrid duplex between DNA and partially methylated nDNA should possess higher specificity than a duplex between DNA with fully methylated nDNA, while retaining binding affinity. However, the application of nDNA for *ex vivo* RNA hybridization at low transcript abundance remains completely unexplored. Here, we determined that partially methylated nDNA (N4 nDNA; with 4 methylated nucleotides) probes inhibited reverse transcription of oncogenic miRNA miR-21 more efficiently than canonical DNA probes or highly methylated nDNA probes (all probes share the same sequence) and with an efficiency rivaling LNA probes. Subsequently, we performed *in situ* hybridization analysis using a miR-21-expressing colorectal cancer cell line (HCT116). HCT116 stained with N4 nDNA probes revealed a greater detection intensity and specificity than HCT116 stained with canonical DNA probes. Consistently, enzyme-linked immunosorbent assays revealed that miRNA hybridization efficiency of N4 nDNA probes was greater than that of canonical DNA probes at cellular transcript levels. Given that N4 nDNA probe is immune-negative and DNase I-resistant, partially methylated nDNA could be further developed to have significant applications in biotechnology and medicine.

## Introduction

**Mi**cro**RNAs** (miRNAs) are a cluster of small, non-coding RNA, generally, 19–24 nucleotides in length, that play essential roles in regulating gene expression by binding to specific target **m**essenger **RNAs** (mRNAs) and inducing mRNA degradation (Shah et al. 2016). Previous studies have shown several correlations between aberrant miRNA expression and various human diseases (Reddy 2015; Shah et al. 2016; Zhang et al. 2007), and as such, typical miRNAs are considered diagnostic and prognostic biomarkers. One particular miRNA, miR-21, has been proposed as a critical diagnostic biomarker for various cancers and is present in multiple tumor tissues (Kriegel et al. 2012; Tang et al. 2014). In particular, human colorectal cancer worldwide is the third leading cause of death amongst all cancer deaths and has a low disease-free survival rate due to typically late-stage diagnosis, shortage of effective therapies, and high metastases rates. Therefore, the development of a technique that can efficiently sense, identify, and locate diagnostic miRNA indicative of malignancy is critical in the diagnosis and treatment of this and other cancers.

Current miRNA detection methods include ***I**n **S**itu* **H**ybridization (ISH) (Nielsen 2012), **R**everse **T**ranscription **q**uantitative **PCR** (RT-qPCR) (Chen et al. 2005), northern blotting (Várallyay et al. 2008), and miRNA microarray profiling technology (Li and Ruan 2009). Of these methods, only ISH provides simultaneous information about miRNA expression level and localization at the single-cell level by utilizing DNA oligonucleotides complementary to specific miRNAs as a detection probe. However, the main challenge of using such DNA as probes is the lack of hybridization specificity and sensitivity, as miRNAs are very short with high homology (many other RNAs could share parts of the same sequence). Moreover, intracellular miRNA levels can be challenging to detect since the ISH output signal scales with the amount of target miRNA. To ameliorate some of these issues, nucleotide analogs such as **L**ocked **N**ucleic **A**cid (LNA) are used as hybridization probes instead of DNA. LNA is unique in that a methylene linkage is included between their 2’-oxygen and 4’-carbon, locking the ribose in the 3’-endo conformation and promoting the formation of A-form duplexes. This is especially key as A-form duplexes are the main conformation that duplex form RNA takes, and RNA binding with LNA probes yields high hybridization affinity due to the stabilization of hydrogen bonds between LNA and miRNA without losing specificity (Kaur et al. 2006). As such, LNA probes in ISH are referred to as the “golden standard” (Jørgensen et al. 2010; Pena et al. 2009; Wienholds et al. 2005). However, this does not preclude possibilities of further optimization of even more efficient ISH probes in the perspective of differentiating high homologous miRNAs.

As such, we sought to investigate alternative non-LNA-based ISH probes. In our previous research, nDNA, a new type of DNA analog containing site-specific methyl phosphotriester (MPTE) inter-nucleoside linkage(s) (**Fig. 1**), was investigated and applied to the DNA sequencing based on field effect transistor (Chen et al. 2013). The advantages of MPTE modifications on the DNA backbone result in higher nuclease stability and the emergence of lipophilic character (Buck 2020). Moreover, MPTE modifications improve duplex hybridization affinity due to the reduction of electrostatic repulsion (reduction of a negative charge for each MPTE linkage) between the strands of an nDNA/DNA,RNA duplex (Chou et al. 2022; Huang et al. 2018; Kuo et al. 2020; Meade et al. 2014; Miller et al. 1986; Moody et al. 1989). Buck, et al. reported that nDNA is capable of hybridizing HIV-1 RNA under *in vitro* conditions and likely is capable of inhibiting HIV-1 infection (Buck et al. 1990). However, the application of nDNA for cellular RNA hybridization at low transcript abundance remains completely unexplored.

**Figure 1.**
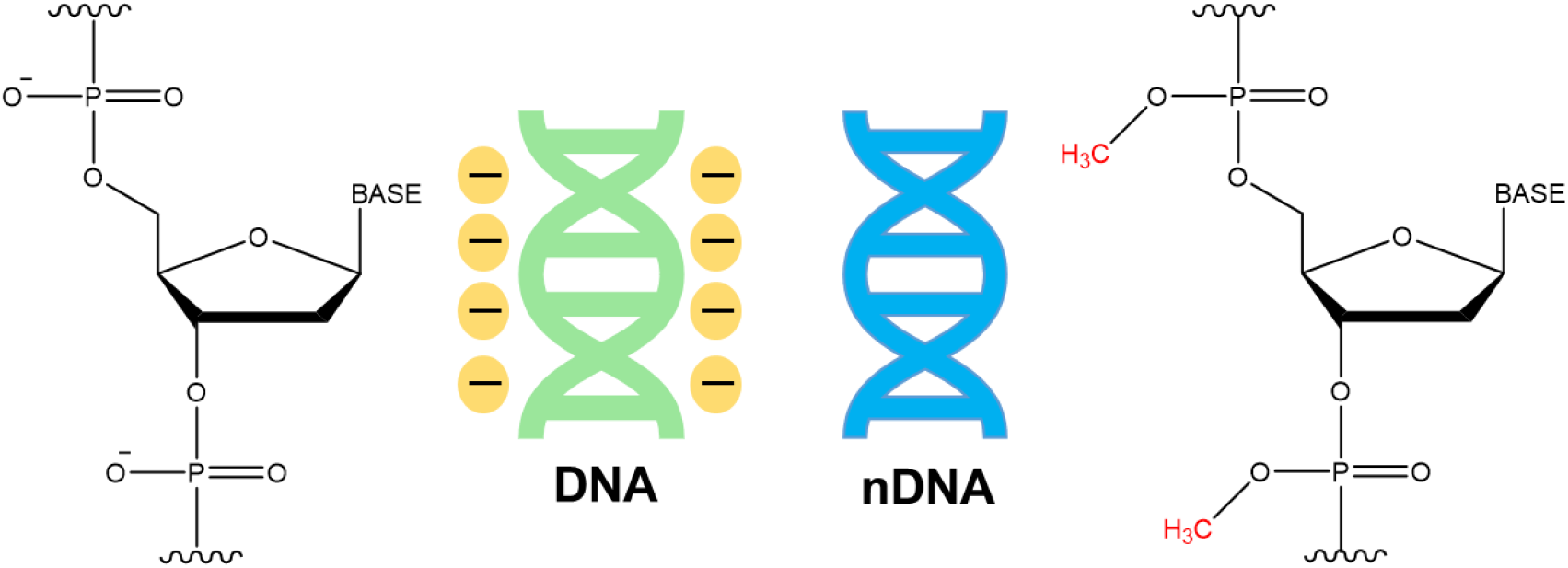
The structure of canonical DNA with a negatively charged phosphodiester backbone and neutralized DNA (nDNA) with a methyl phosphotriester backbone, respectively.

From a thermodynamic perspective due to steric effects, a hybrid duplex between DNA and partially methylated nDNA should possess higher specificity than a duplex between DNA with fully methylated nDNA, while retaining binding affinity. Because of the increased hybridization specificity and stability of partially methylated nDNA strands and the latest advances in synthetic techniques that allow more control over MPTE design, we applied nDNA to ISH studies to detect various endogenous miRNA (miR-21) as a proof of principle. The study showed that because of the improved probe hybridization stability, without losing probe specificity, nDNA ISH probes could be used as a miRNA detection technique for diagnostic miRNA expression under *ex vivo* conditions.

### Materials and Methods

All of the chemicals were purchased from Sigma-Aldrich, Inc. (St. Louis, MO, USA) unless specified otherwise.

#### Cell culture

The human colorectal cancer cell line HCT116 was kindly provided by Professor Li-Jen Su from the Institute of Systems Biology and Bioinformatics, National Central University, Taipei, Taiwan(Lai et al. 2022). HCT116 cells were cultured with McCoy’s 5A Medium (GIBCO, Invitrogen, Waltham, USA) supplemented with 10% fetal bovine serum (FBS; GIBCO, Invitrogen) and 1% each of penicillin and streptomycin (GIBCO, Invitrogen) in a humidified incubator (Thermo Fisher Scientific Waltham, USA) under 95% (v/v) air and 5% C (v/v) at 37°C.

#### miRNA hybridization

The hybridization reaction containing miR-21 (0.1 nM) (mdbio, Taipei, Taiwan); 1, 2, 4, or 16 nM of DNA/nDNA/LNA probe; 1X SSC buffer (BioChain Institute Inc., Newark, USA); and UltraPure Distilled Water (Invitrogen) was incubated at 37°C for 30 min. A “scramble” random probe sequence (negative control; **Table 1**) was generated to validate the specific inhibition of miR-21 RT by antisense oligonucleotide probes due to their sequence specificity but not due to non-specific hybridization.

**Table 1.**
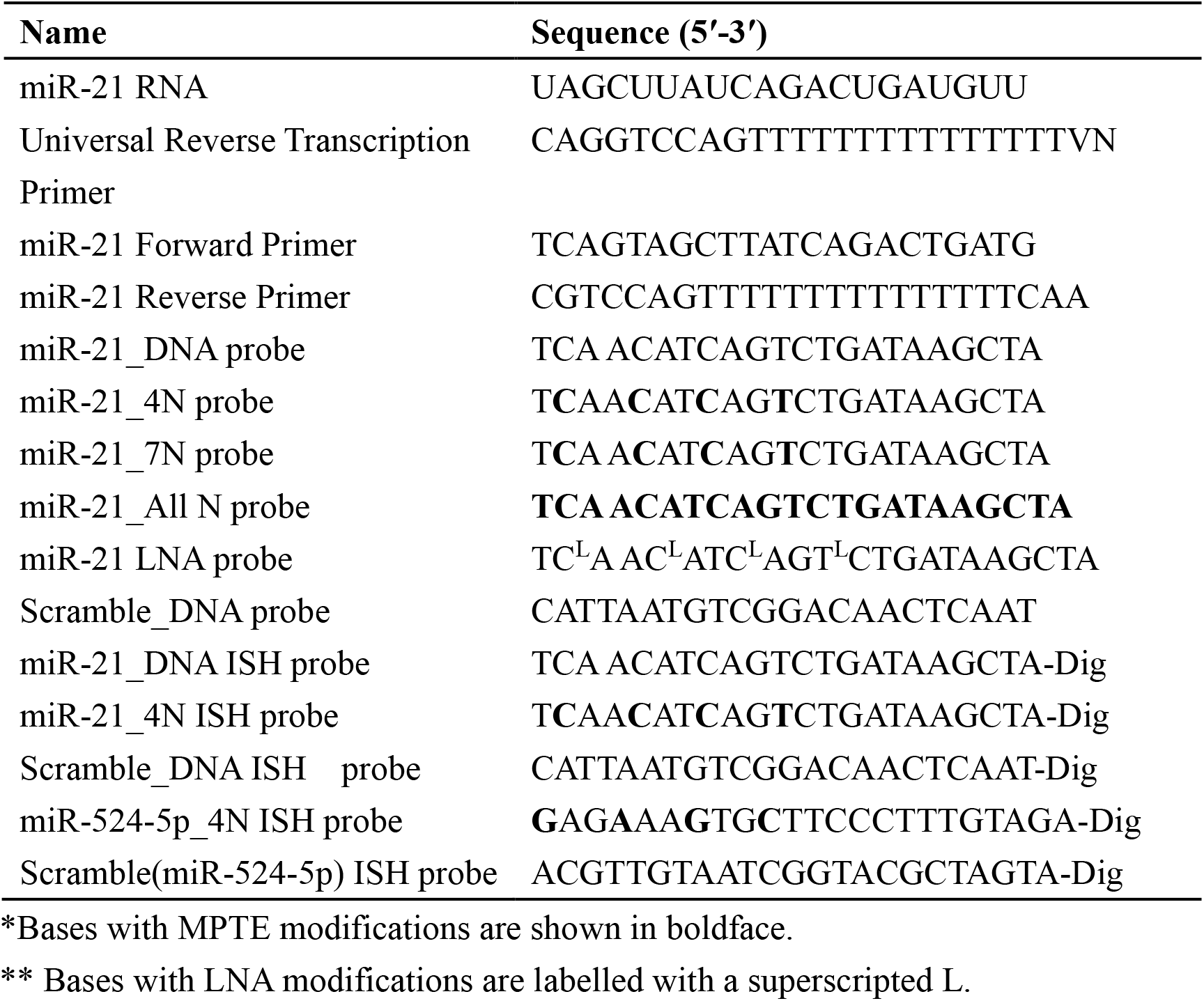
Oligonucleotides sequences used in the study.

#### Reverse transcription (RT)

miRNA (1 nM) was converted into cDNA using adenosine 5′-triphosphate (1 mM) (**N**ew **E**ngland **B**iolabs (NEB), Ipswitch, MA, USA), deoxynucleotide (dNTP) solution mix (NEB), 10X Poly(A)Pol reaction buffer (NEB), SuperScript® III reverse transcriptase (Invitrogen) (10 U), *E. coli* Poly(A) polymerase (NEB) (10 U), RNase inhibitor (0.4 U) (Thermo Fisher Scientific), UltraPure distilled water (Invitrogen), and universal RT primer (mdbio) (1 μM) to a final volume of 10.2 μL. RT was performed using an initial step at 42°C on a water bath for 15, 30, 45, and 60 min, respectively, then kept on ice until further use.

#### Real-time quantitative PCR (qPCR)

Expression of miRNA was determined by TaqMan® MicroRNA Assays using an SYBR Green Master Mix (Applied Biosystems, Foster City, USA) on a StepOne™ Real-Time PCR System (Applied Biosystems) as described previously (Li et al. 2019). Three replicates were analyzed through StepOne™ software v2.3. The qPCR reactions (10 μL) contained SYBR Green Master Mix (5 μL) (Applied Biosystems), forward/reverse primers (1 μL each) (mdbio), and ultrapure distilled water (2 μL) (Invitrogen). qPCR was performed using an initial step at 95°C for 5 min for DNA denaturation, followed by 40 cycles of 95°C for 10 s (DNA denaturation), and 60°C for 15 s (primer annealing and elongation). For a fair comparison, all Ct values of qPCR experimental trials in this study were calculated using a unifying threshold.

#### Methyl phosphotriester (MPTE) modified neutralized DNA (nDNA) probes

Oligonucleotide primers and nDNA probes containing MPTE were purchased from Helios Biotech Inc., Taipei, Taiwan. A probe to test for miR-21 with the sequence 5′-TCAACATCAGTCTGATAAGCTA-3′ with three MPTE modifications spaced every three nucleotides was analyzed by high-performance liquid chromatography (HPLC) and Liquid Chromatography–Mass Spectrometry to confirm the purity and molecular weight following synthesis (**Figs. S1 and S2**).

#### In situ hybridization (ISH)

*In situ* hybridization was performed following the protocols established previously with some modifications (Erener et al. 2021). Briefly, cells were cultured on an 8-well plastic plate (ibidi, Gräfelfing, Germany). After the existence of 80% confluent cells, the cells were fixed in 4% paraformaldehyde (Thermo Fisher Scientific) for 20 min at room temperature. Cells were then digested using proteinase K (15 mg/L) (Thermo Fisher Scientific) at room temperature for 8 min, washed thrice with phosphate buffer saline (PBS), and then fixed in 4% paraformaldehyde for 15 min again. Cells were then pre-hybridized with pre-hybridization buffer (BioChain Institute Inc.) for 4 h at 50°C. Cells were then hybridized with a hybridization buffer (BioChain Institute Inc.) containing the hybridization probe for 8 h at 40°C. After hybridization, cells were rinsed with 2-fold saline-sodium citrate buffer (SSC) (BioChain Institute Inc.) at 37°C for 5 min; then rinsed with 1.5-fold SSC at 37°C for 5 min, and finally rinsed with 0.5-fold SSC at 45°C for 5 min. This procedure was repeated twice. Cells were blocked with blocking agents (BioChain Institute Inc.) for 1 h at room temperature. Anti-DIG antibody conjugated to alkaline phosphatase (BioChain Institute Inc.) at 1:200 dilution in PBS was applied to the cells for 4 h at room temperature. The cells were then finally visualized using a combination of NBT (nitro-blue tetrazolium chloride) and BCIP (5-bromo-4-chloro-3’-indolyphosphate p-toluidine salt) (**Fig. S3**) on an IX83 inverted microscope under 40X amplification without oil (Olympus, Tokyo, Japan).

#### Enzyme-linked immunosorbent assay (ELISA)

The enzyme-linked immunosorbent assay was performed following the protocols established previously with some modifications (Kinman and Pompano 2019). Briefly, cells were cultured on a 24-well microplate (Falcon, Corning, Glendale, USA). After the existence of 80% confluent cells, the cells were fixed in 4% paraformaldehyde (Thermo Fisher Scientific) for 20 min at room temperature. Cells were then digested using proteinase K (15 mg/L) (Thermo Fisher Scientific) at room temperature for 8 min, washed thrice with PBS, and then fixed in 4% (w/v) paraformaldehyde for 15 min again. Cells were then pre-hybridized with pre-hybridization buffer (BioChain Institute Inc.) for 4 h at 50°C. Cells were then hybridized with a hybridization buffer (BioChain Institute Inc.) containing a hybridization probe for 17 h at 40°C. After hybridization, cells were rinsed with 2-fold SSC (BioChain Institute Inc.) at 37°C for 5 min; then rinsed with 1.5X SSC at 37°C, for 5 min; and finally rinsed with 0.5-fold SSC at 45°C for 5 min. This procedure was repeated twice. Cells were blocked with blocking agents for 1 h at room temperature. Anti-DIG antibody conjugated to alkaline phosphatase (BioChain Institute Inc., USA) at 1:200 dilution in phosphate buffer saline was applied to the cells for 4 h at room temperature. The cells were then finally visualized using a combination of NBT and BCIP. The supernatant of each sample was also moved to a 96-well microplate (Falcon, Corning) at 8, 10, 12, and 14 h, and absorbance values of the respective supernatants were read at 560 nm in a SpectraMax iD3 microplate reader (Molecular Devices, Downingtown, USA).

#### Oligonucleotide 3′ labeling

Oligonucleotide 3′ labeling was performed using the DIG Oligonucleotide 3′-End Labeling Kit, 2^nd^ generation (Roche, Basel, Switzerland). Each indicated oligonucleotide (100 pmol) was added to each reaction vial along with UltraPure distilled water (Invitrogen) to a final volume of 10 µL. The reaction vials were placed on ice, and to the reaction vial we added 5-fold reaction buffer, 1-fold CoCl2 solution (0.5 mM) (Roche), DIG-2’,3’-dideoxyuridine-5’-triphosphate (ddUTP) solution (5 mM) (Roche), and terminal transferase (50 μM) (Roche). The reagents were mixed thoroughly *via* pipetting and were incubated at 37°C for 15 min. Reactions were stopped by adding 2 µL EDTA (0.2 M; pH 8) (Protech, Taipei, Taiwan).

#### Innate immune response assay

An innate immune response assay was performed following established protocols described previously (Yi et al. 2013). Briefly, THP1-Dual™ KI-hSTING-R232, NF-κB-SEAP, and IRF-Lucia reporter monocytes (InvivoGen, Hong Kong, SAR, China) were seeded in a 96-well plate at a density of ∼5.0 × 10^5^/mL. Cells were treated with cGAMP (positive control), DNA probe, or N4 nDNA probe (10 μg/mL and 100 μg/mL) (sequences available in **Table 1**). After 24 h, cell supernatants were harvested, and the luciferase activities were determined using QUANTI-Luc™, a Lucia™ detection reagent.

## Results and Discussion

### Antisense-inhibition of miRNA RT by N4 nDNA (four MPTE linkages) in synthetic human plasma

In principle, a single strand RNA molecule is required for a RT reaction. Antisense oligonucleotide probes (DNA) are used to hybridize RNA strands and thus can inhibit the RT reaction, which can be quantified *via* quantitative PCR (qPCR); the greater the number of cycle threshold (Ct) required for successful RT-qPCR, the greater the inhibitory characteristics of the probe (**Fig. 2A**). Previously, it has been predicted that antisense oligonucleotides containing a low number of methylated linkages may be more thermodynamically stable binders than canonical and highly-methylated oligonucleotides due to steric effects (Kuo et al. 2020) (**Fig. S4**). Thus, to investigate the efficacy of nDNA as an antisense oligonucleotide probe, we first determined the ability of a partially methylated nDNA with four MPTE linkages (N4 nDNA) to specifically inhibit RT of an oncogenic miRNA miR-21, and compared the efficacy to the canonical DNA, LNA, and nDNA with more methylated linkages (N7 nDNA and All_N nDNA, seven and all MPTE linkages, respectively) of identical sequence; all sequences can be found in **Table 1**. Given that somatic adult cell volume is ∼5,000 μm^3^(Cohen and Studzinski 1967) and that intracellular miR-21 copy numbers range between ∼700–12,000 molecules per cell (Lim et al. 2003), the intracellular miR-21 concentration would be ∼0.2–4 nM (See **Table S1** for detailed calculation). Therefore, to estimate the sensitivity and efficacy of antisense nDNA probes in simulated cellular environments, we performed the miR-21 RT reactions in synthetic human plasma with a miR-21 concentration of 0.1 nM, which is slightly below the lower range of intracellular miR-21 concentration and would show that our method is sensitive enough for real cellular measurements.

**Figure 2.**
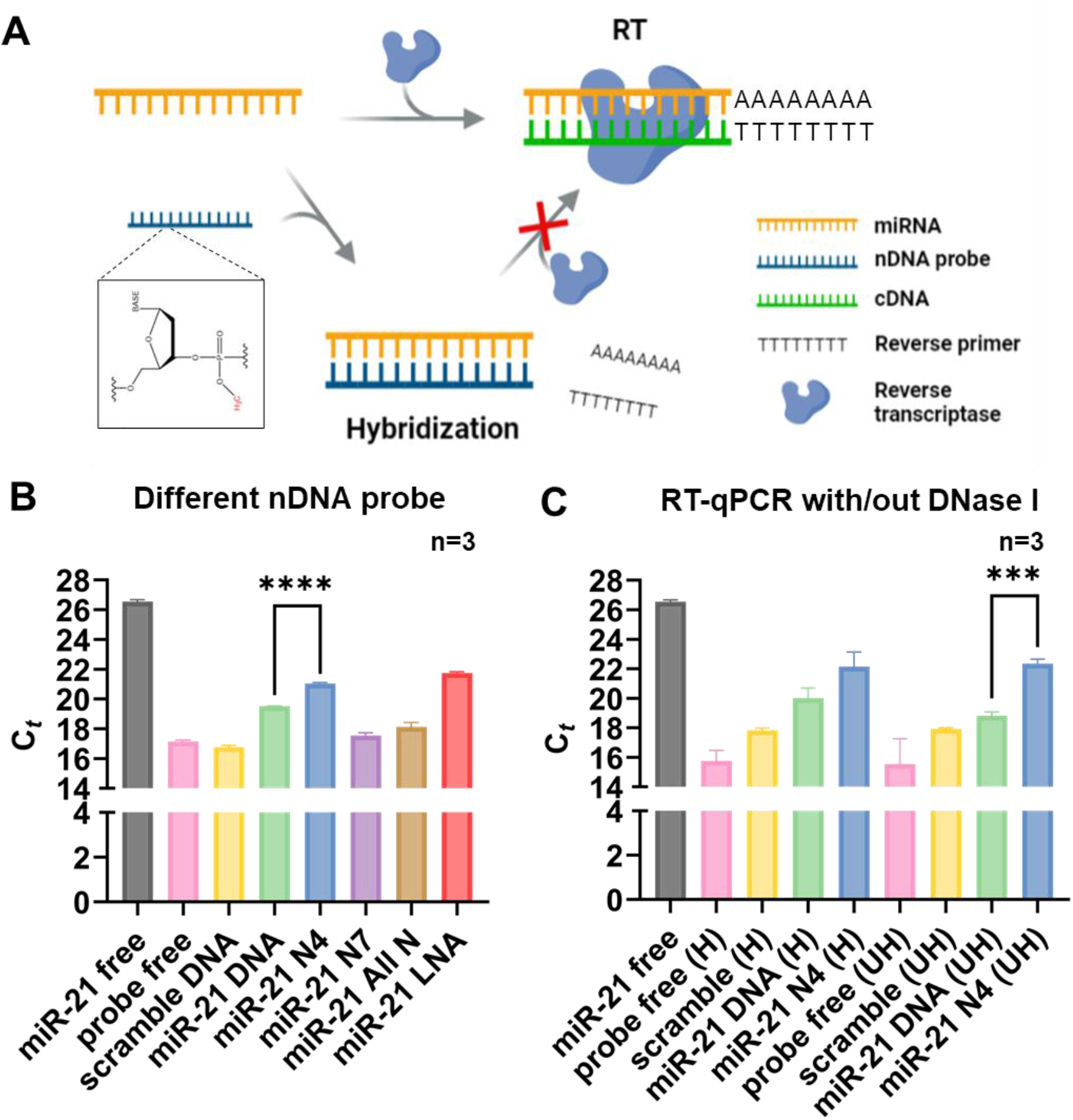
Sensitive and specific inhibition of miR-21 reverse transcription (RT) by an N4 antisense nDNA probe in synthetic human plasma. **(A)** Schematics of the inhibition of miRNA RT due to formation of the miRNA/nDNA duplex. **(B)** The inhibition of miR-21 RT by different antisense oligonucleotide probes. The inhibition efficacy of antisense oligonucleotide probes is measured using qPCR. **(C)** DNase I resistance of the N4 nDNA antisense oligonucleotide probe was revealed by RT-qPCR with denatured DNase I (H, heated) or with functional DNase I (UH, unheated). In all experimental trials, scramble DNA, canonical DNA, and LNA probes are used to compare the efficacy of nDNA probes. The statistical significance was verified by an unpaired t-test, *: p< 0.05; **: p< 0.01; ***: p< 0.005; ****: p< 0.001. The bars represent the standard errors and the symbols represent the mean from three independent experimental replicates.

Compared to the RT reactions free of antisense oligonucleotide probes (positive control), the N4 nDNA probe more effectively inhibited the miR-21 RT reaction compared to equal concentration (4 nM) of canonical DNA probe, highly methylated nDNA probes (N7 and All_N), and a negative control “scramble” probe with random sequence, confirming our initial hypothesis (**Fig. 2B**). Moreover, the N4 nDNA probe inhibited the miR-21 RT reactions with an efficiency comparable to that of the corresponding LNA probe, the theoretical “gold standard” of currently used antisense probes (**Fig. 2B**). Therefore, our data suggest that the inhibition of the RT reactions by the N4 nDNA probe is efficient and specific (due to miRNA/N4 nDNA hybridization), and we thus used only the N4 nDNA probe rather than other nDNA probes for subsequent experiments due to its greater efficacy.

One of the reasons nDNA probes could have more RT inhibition efficacy than traditional DNA probes in cells is that the MPTE backbone may inherently be resistant to DNase degradation, which may be present significantly in standard biological samples. Since the polyA tail is a well-known exonuclease inhibitor (Slomovic et al. 2005) and is also an essential component to initiate the RT reaction, we only tested the resistance of nDNA against endonuclease DNase I. As such, we next quantified the efficacy of RT inhibition by both canonical DNA and N4 nDNA probes in the presence and absence of functional DNAse I by performing qPCR assays under both DNase I-heated (H) and DNase I-unheated (UH) conditions (**Fig. 2C**). Heating of DNase I would result in protein denaturation and loss-of-function, while unheated functional DNase I would hydrolyze DNA/nDNA and thus may abolish the inhibition of RT by the probes. While in both the presence and the absence of functional DNase I both canonical DNA and N4 nDNA probes were significantly more functional than in probe-free conditions (p< 0.001 in heated conditions, p=0.0018 (DNA) and p= 0.0009 (N4 nDNA), **Table S2**), functional DNase I (*i*.*e*., the unheated conditions) did not inhibit the efficacy of an N4 nDNA probe compared to the heated conditions, and N4 nDNA probes were significantly more effective at inhibition of miR-21 RT in the presence of functional DNase I compared to canonical DNA probes (p= 0.0001) (**Fig. 2C, Table S2**). Moreover, the addition of heated DNase I in the RT reactions did not affect the efficacy of N4 nDNA probe inhibition of miR-21 RT (as measured by qPCR) in the probe-free experimental trials (**Fig. 2C**). These suggest that N4 nDNA could withstand potential DNase hydrolysis in simulated cellular environments and thus could be a more competent candidate than canonical DNA for cellular or physiological applications using antisense oligonucleotide probes.

We also observed that the N4 nDNA probe inhibited the miR-21 RT reaction with more efficacy than the canonical DNA probe at short incubation times before qPCR (15 minutes; common RT incubation time is ∼60 minutes) (**Fig. 3A**). However, at longer incubation times before qPCR (30 minutes), the nDNA probe and the canonical DNA probe appeared to inhibit the RT reaction of miR-21 with similar efficacy (but the efficacy of both was still stronger than a scramble sequence probe). The discrepancy in probe effectiveness with variable incubation times before qPCR may be due to the fact that the hybridization kinetics rates of the various probes with miR-21 are variable, and that N4 nDNA may hybridize with miR-21 faster than the DNA probe at the concentrations used in this study. However, longer incubation times (longer than 15 minutes) before qPCR allow all of the probes to hybridize fully to miR-21, counteracting the apparent initial N4 nDNA inhibitory advantage. Nevertheless, decreasing the required incubation time before qPCR analysis by using N4 nDNA (to 15 minutes, which is less than half of the traditional incubation time used for similar studies) shows its usefulness in increasing the throughput of such studies, and subsequent qPCR analyses use this incubation time.

**Figure 3.**
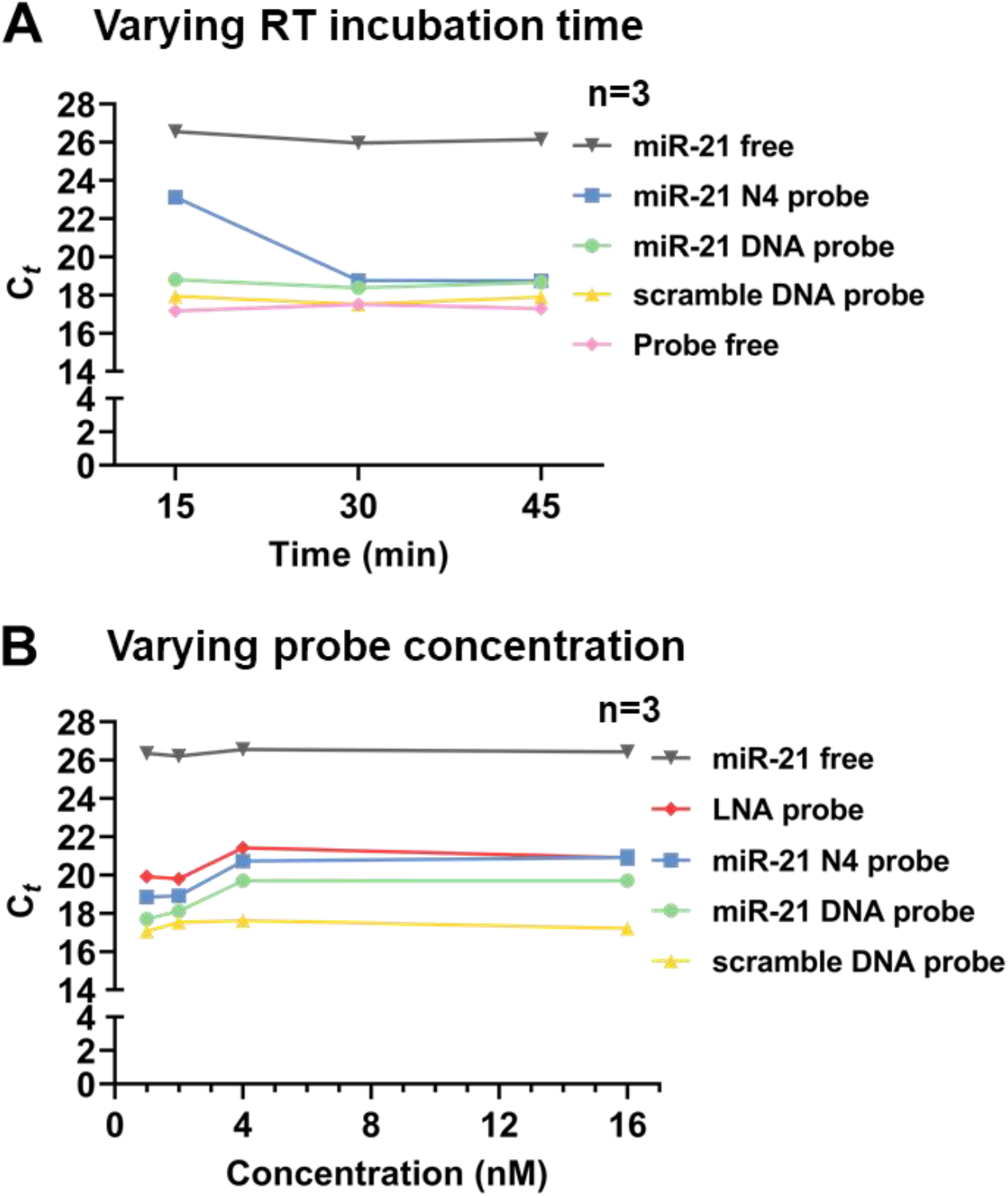
The specific inhibition efficiency of miR-21 reverse transcription (RT) by an antisense N4 nDNA probe in synthetic human plasma. **(A-B)** The inhibition efficiency of antisense oligonucleotide probes for miR-21 RT with varying RT incubation times before qPCR **(A)** and varying probe concentrations **(B)**. The bars represent the standard errors and the symbols represent the mean from three independent experimental replicates.

Next, we managed to determine whether the amount of probe present could modulate the efficacy of miR-21 RT inhibition by assaying qPCR studies with probe concentrations increasing from 1–16 nM for canonical DNA, N4 nDNA, LNA, and a scramble sequence probe (with a 15-minute-incubation before qPCR) (**Fig. 3B**). It appeared that for canonical DNA, N4 nDNA, and LNA probes, their RT inhibition efficacy increased for concentrations up until 4 nM (a miRNA-to-probe ratio of 1 to 40), but the efficacy reached a plateau beyond this concentration and 16 nM probe concentration resulted in no significant difference in inhibition compared to 4 nM probe concentration for all probes (**Fig. 3B**). However, in all concentrations tested, the N4 nDNA probe appeared to be more effective than the canonical DNA probe, suggesting their superiority for RT inhibition. Moreover, the N4 nDNA probe is as effective as the LNA probe when the probe concentration is above 4 nM, but is slightly less effective (ΔCt <1) than the LNA probe below 4 nM. Once again, all specific probes were more effective in miR-21 RT inhibition than the scramble probe.

### Quantification of miRNA ISH by N4 nDNA in cancer cell HCT116

The fact that the N4 nDNA probe appeared to be more effective at miR-21 RT inhibition than the canonical DNA probe under certain conditions gives credence to the hypothesis that an N4 nDNA oligonucleotide could bind more strongly to miR-21 than a canonical DNA probe at the given conditions. Thus, we next quantified the efficacy of N4 nDNA as a probe for ISH in an *ex vivo* colorectal cancer cell line (HCT116) expressing miR-21 (**Fig. 4**). We designed N4 nDNA, canonical DNA, and scramble DNA probes for miR-21 each with a digoxigenin (Dig) tag. These Dig-tagged antisense probes were then applied to the HCT116 cell culture, and presumably, due to the high concentration of miR-21 within the cells (∼12,000 copies per cell, or 4 nM) (**Table S1**), the probes would be uptaken, and hybridized to intracellular miR-21. The addition of an anti-Dig antibody conjugated to alkaline phosphatase was then applied to the cell culture, and the magenta stain (due to a combination of nitro-blue tetrazolium (BNT) and bromo-chloro-indolyphosphate (BCIP)) was used to visualize the location of the probe in comparison with the cells. While a scramble sequence showed no localization to HCT116 cells due to its inability to hybridize to miR-21, the miR-21 antisense DNA probe appeared to localize to some degree within HCT116 cells, showing its ability to perform basic cellular ISH (**Fig. 4**). Remarkably, the miR-21 antisense N4 nDNA probe significantly localized to the HCT116 cells and we were able to visualize the cells with high contrast, suggesting that N4 nDNA probes could potentially be more effective than canonical DNA probes for cellular ISH. Given that miR-21 is an intrinsic miRNA in HCT116 cells, we also managed to use an exogenous miRNA not present in the HCT116 cell line to further examine the application of antisense N4 nDNA probes to perform ISH. Therefore, synthetic miR-524-5p was introduced to the HCT116 cells *via* transfection, and the ISH trials suggested that the miR-524-5p antisense N4 nDNA probe significantly localized to the HCT116 cells, while a scramble sequence showed no localization to the miR-524-5p-transfected HCT116 cells (**Fig. S5**). Together, our data validated the fact that partially methylated nDNA can serve as a sensitive and specific antisense oligonucleotide probe for cellular ISH of both intrinsic and exogenous miRNA.

**Figure 4.**
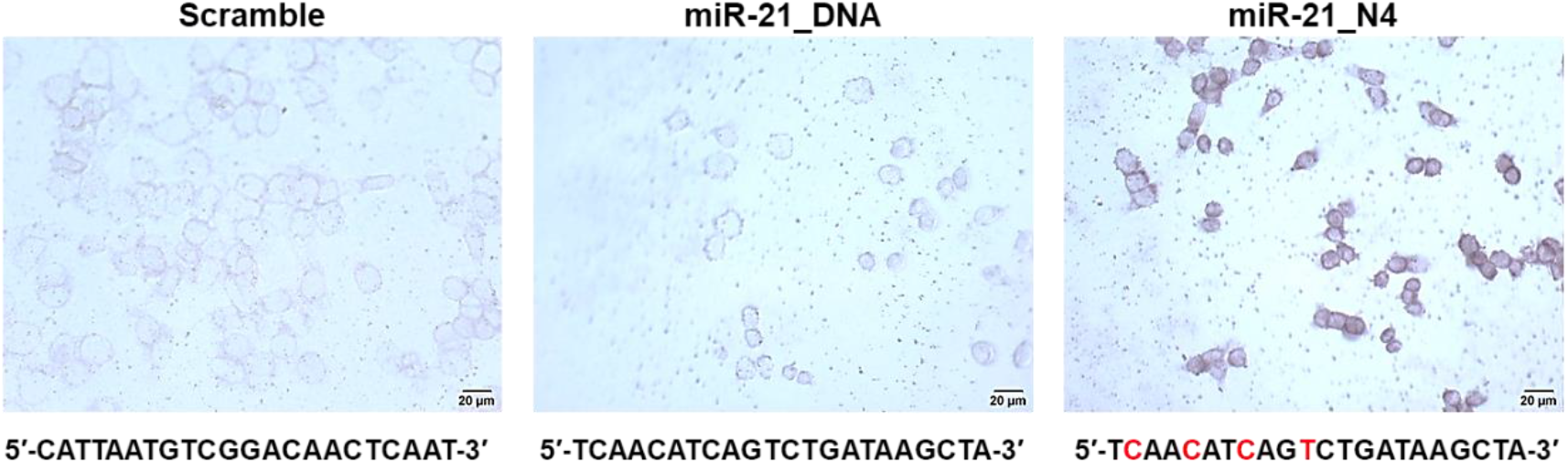
*In situ* hybridization (ISH) of miR-21 in the HCT116 cells. ISH images of cells stained using the scramble oligonucleotide, antisense canonical DNA probe, and antisense N4 nDNA probe, respectively. The methylated nucleotides on the N4 nDNA probes are highlighted in red.

Thus, we next set out to quantify the degree to which N4 nDNA probes may be more effective than canonical DNA probes through an enzyme-linked immunosorbent assay (ELISA) measuring miR-21 expression within HCT116 cells in real-time. The identical Dig-linked antisense probes used in **Fig. 4** were each applied to the HCT116 cell culture in a microplate for the ELISA (**Fig. 5A**), and the miR-21 expression level was tracked over time by measuring the saturation concentration (*i*.*e*., magenta color intensity) of the sample at specific time points. The saturation concentration of both the N4 nDNA and canonical DNA probes increased over time, reaching around 75% by 14 hours after initiation (**Fig. 5B**). This suggested that miR-21 was successfully expressed during this period to some extent. However, it appeared that at intermediate time points (8 and 10 hours), the N4 nDNA probe sample exhibited significantly greater color intensity (∼2-fold) than the canonical DNA probe sample (also shown directly by a photograph of the respective sample wells), which suggests its greater cell membrane permeability and greater ability to hybridize to nascent miR-21. However, at longer time points (12 and 14 hours), the saturation concentration of the N4 nDNA probe sample may not have been significantly higher than the canonical DNA probe sample (although it could have been ∼10% greater), suggesting that the advantage of N4 nDNA may be more pronounced at shorter or intermediate reaction times. The color of the culture with the scramble probe did not appear to change over time, despite the fact that miR-21 was confirmed to have been expressed during the course of the reaction.

**Figure 5.**
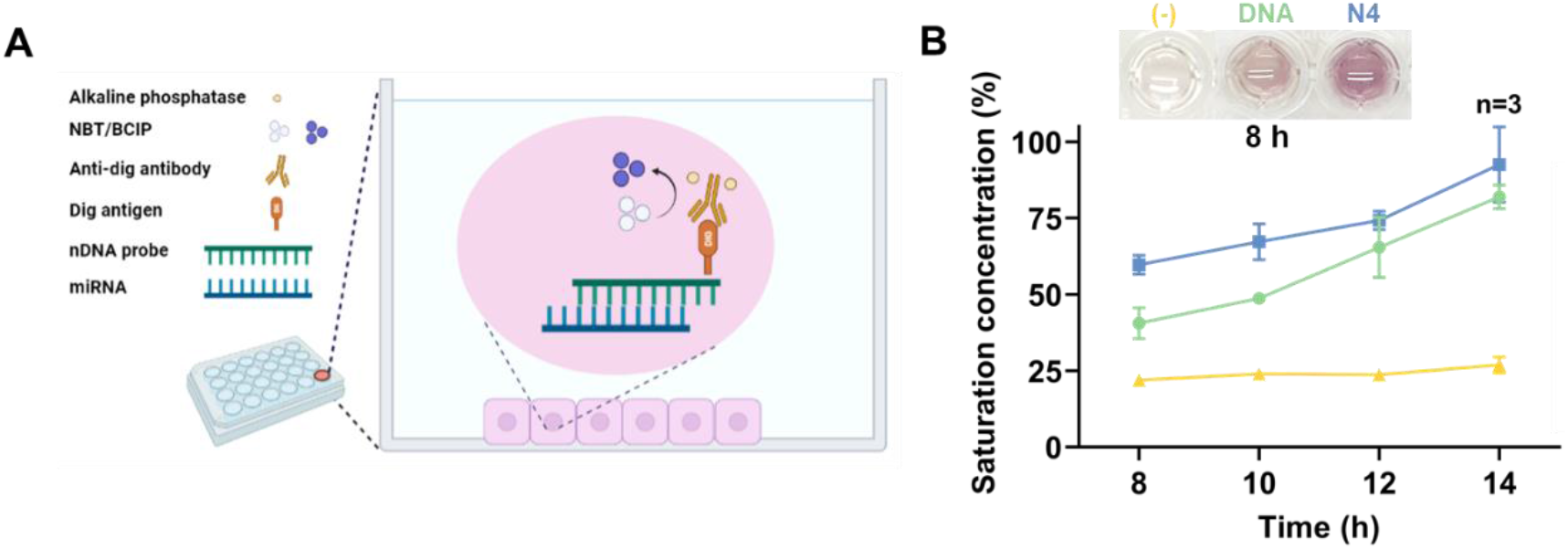
An enzyme-linked immunosorbent (ELISA) assay for measurement of miR-21 miRNA expression in HCT116 cells using antisense oligonucleotide probes. **(A)** Schematics of the microplate-based ELISA assay with digoxigenin (Dig)-linked antisense oligonucleotide probes for MiR-21 miRNA hybridization. An anti-Dig antibody conjugated to alkaline phosphatase is used to visualize cells (magenta color) using a combination of nitro-blue tetrazolium (BNT) and bromo-chloro-indolyphosphate (BCIP). **(B)** Time-dependent measurements of the concentration of the BCIP-derived product in the ELISA assay hybridized with either the scramble probe, DNA probe, or N4 nDNA probe. The bars represent the standard errors and the symbols represent the mean from three independent experimental replicates.

## Conclusions

Here, we characterized a new type of nucleic acid analog probe, MPTE-based nucleic acid (*i*.*e*., nDNA), for miRNA expression inhibition and ISH. Moreover, partially methylated nDNA probes were observed to exhibit stronger hybridization affinity and stability than canonical DNA probes, while performing as well as an LNA probe and performing better than canonical DNA probes in inhibition of miRNA expression (especially at shorter timescales). It is postulated that this difference in efficacy may be due to increases in hybridization affinity and stability caused by the reduction of electrostatic repulsion, increases in the hydrophobicity, and/or steric of the phosphate backbone upon methylation. Despite the increase in hybridization affinity and stability, the nDNA binding specificity was unaffected through careful design. While this study focused on ISH, the antisense nDNA probes linked with fluorophore could also be used for *in vitro* oncogene diagnostic assays (Park et al. 2022) and **F**luorescent ***I**n **S**itu* **H**ybridization (FISH). Given that the number of MPTE modifications in an nDNA probe can be tuned finely, it is plausible that gene expression level can also be tuned based on the associated changes in hybridization affinity upon methylation, but not based on a dose-dependent manner. Thus, as N4 nDNA probes do not elicit a strong immune response in cells (**Fig. S6**) and are resistant to DNase I (**Fig. 2C**), future development and optimization of nDNA probes for *in vivo* gene expression modulation could be the way forward for personalized genetic medicines using the established antisense oligonucleotide sequences targeting carcinogenic miRNA and mRNA (Moitra et al. 2022).

## ACKNOWLEDGMENTS

This study is supported by the Ministry of Science and Technology of Taiwan (MOST) (110-2222-E-008-002-MY2; 108-2221-E-008-056-MY3). Tony Z. Jia is supported by the Japan Society for the Promotion of Science (JSPS) (Grants-in-aid 21K14746). Po-Hsiang Wang is also supported by the Research and Development Office as well as Research Center for Sustainable Environmental Technology, National Central University, Taiwan.

## Author contributions

W.Y.C. and P.H.W. conceptualized this study. C.W.C. and P.H.W. performed the RT-qPCR, ISH, and ELISA experiments. Y.H.C. performed the cell-based immune response assays. W.Y.C, T.Z.J., B.C.M.T., T.Z.J., and P.H.W. analyzed the data. P.H.W., W.Y.C., and T.Z.J. wrote this manuscript with assistance from all the authors. All the authors were participating in drawing the figures and tables.

