## Supplemental information for "Sensitive and specific miRNA *in situ* hybridization using partially methylated phosphotriester antisense DNA probes"

### Table of Contents


**Supplemental Methods**

**Innate immune response assay**

**Transient transfection of miRNA Mimics**

**Supplemental Figures**

**Fig. S1.** High-performance liquid chromatography (HPLC) of antisense N4 nDNA probe.

**Fig. S2.** Mass spectrometry to confirm the purity and molecular weight of antisense N4 nDNA probe.

**Fig. S3.** Reaction mechanism of BCIP/ NBT substrate with alkaline phosphatase.

**Fig. S4.** Hybridization efficiency of different DNA and nDNA probes at varying concentrations.

**Fig. S5.** *In situ* hybridization (ISH) of miRNA miR-524-5p in HCT116 cells.

**Fig. S6**. Innate immune response of canonical DNA and antisense N4 nDNA probes

**Supplemental Tables**

**Table S1.** Intracellular miR-21 concentration

**Table S2.** p-value of RT-qPCR with/out DNase I

**Appendix**

**References**

#### Supplemental Methods

**Innate immune response assay.** Innate immune response assay was performed following established protocols described previously [1]. Briefly, THP1-Dual™ KI-hSTING-R232 NF-κB-SEAP and IRF-Lucia reporter monocytes (InvivoGen, HongKong) were seeded in a 96-well plate at a density of ~5.0 × $10^5$/mL. Cells were treated with cGAMP (positive control), DNA probe, or N4 nDNA probe (10 μg/mL or 100 μg/mL) (sequences available in **Table 1**). After 24 h, cell supernatants were harvested, and the luciferase activities were determined using QUANTI-Luc™, a Lucia™ detection reagent.

**Transient Transfection of miRNA Mimics.** The HCT116 cells were transfected with has-miR-524-5p microRNA Mimics (Exiqon, Denmark). Briefly, the HCT116 cells were grown overnight and then transfected with has-miR-524-5p microRNA Mimics (100 nM) using the TransIT-X2 system (Mirus Bio, USA) according to the manufacturer's protocol. The has-miR-524-5p microRNA Mimics (5.6 μL) and TransIT-X2 transfection reagent (1.5 μL) were added to the Opti-MEM™ I Reduced Serum Medium (50 μL) (GIBCO, Invitrogen, USA), followed by an incubation for 30 min at room temperature. The resulting complex was added to each well containing the HCT116 cells in an 8-well plate, and the plate was gently swirled while adding the complex to spread entirely in all parts of the plate. The plate was incubated for 24 h.

#### Supplemental Figures

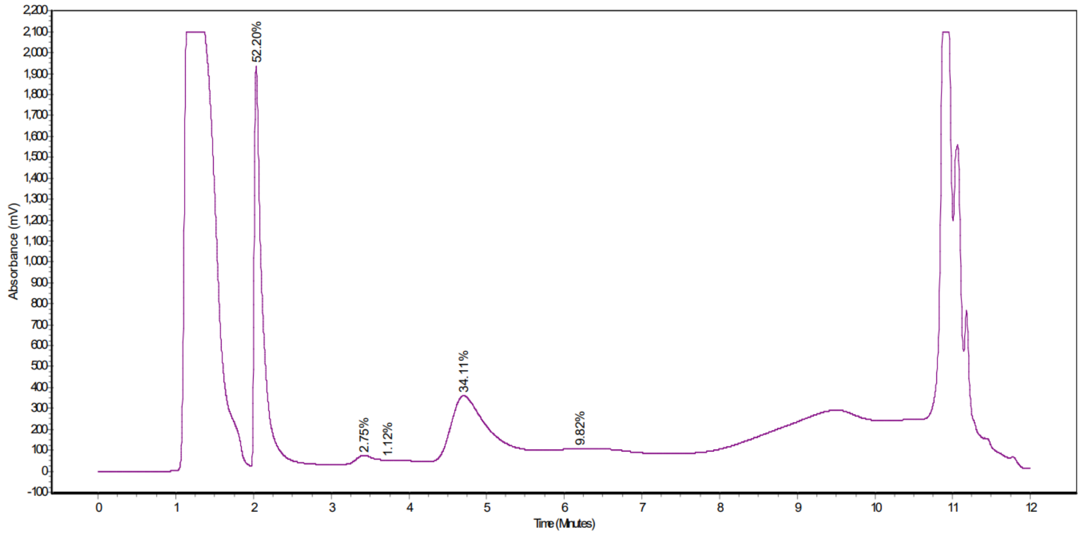


**Fig. S1.** High-performance liquid chromatography (HPLC) of antisense N4 nDNA probe.

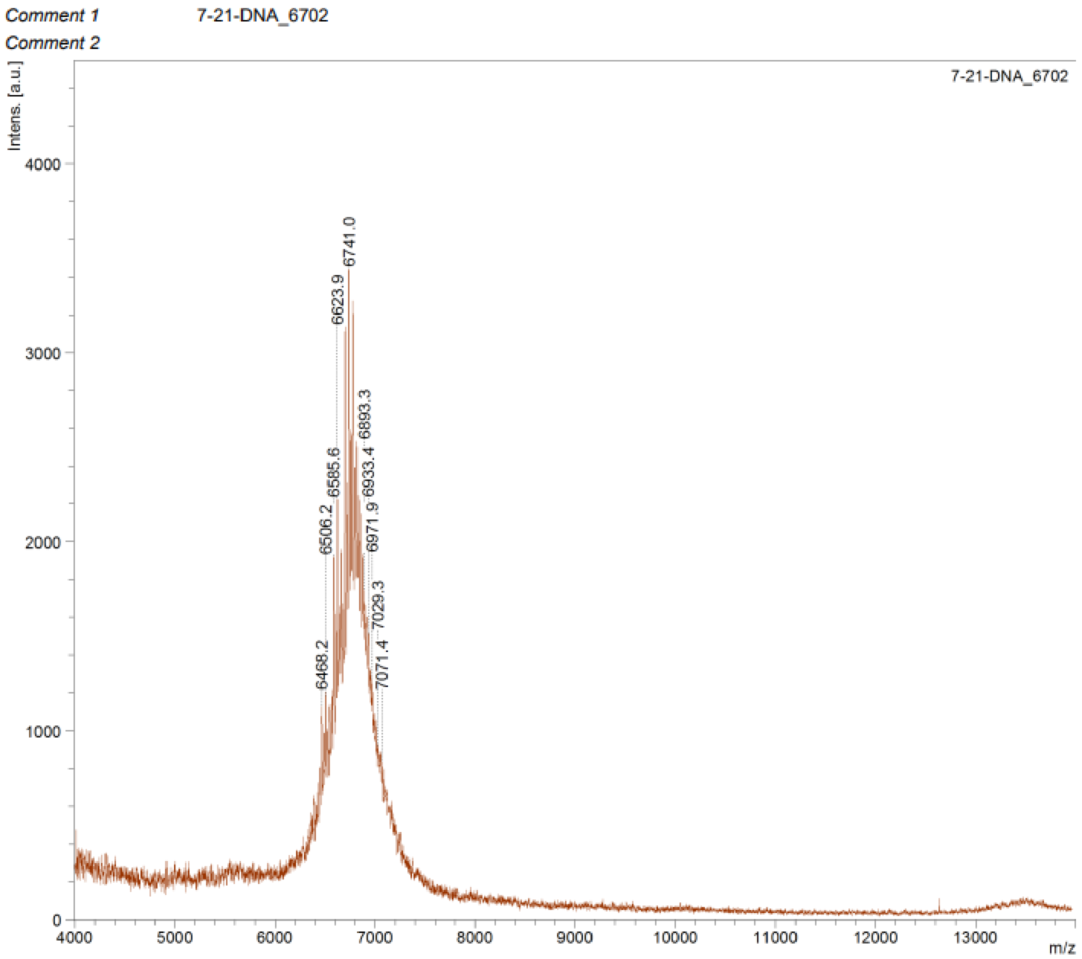


**Fig. S2.** Mass spectrometry to confirm the purity and molecular weight of antisense N4 nDNA probe.

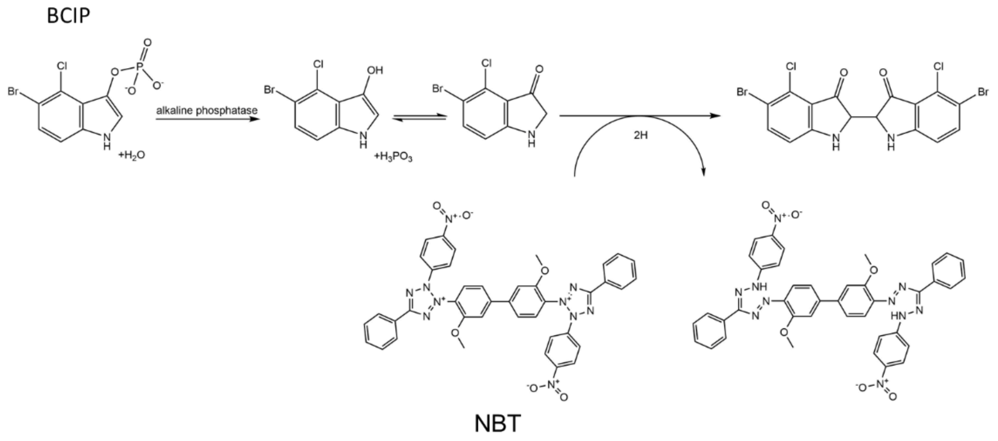


**Fig. S3.** Reaction mechanism of BCIP/ NBT substrate with alkaline phosphatase.

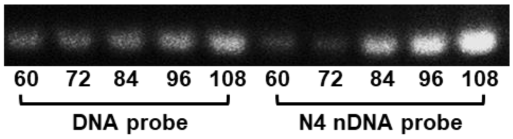


**Fig. S4.** Hybridization efficiency of different DNA and nDNA probes at varying concentrations.

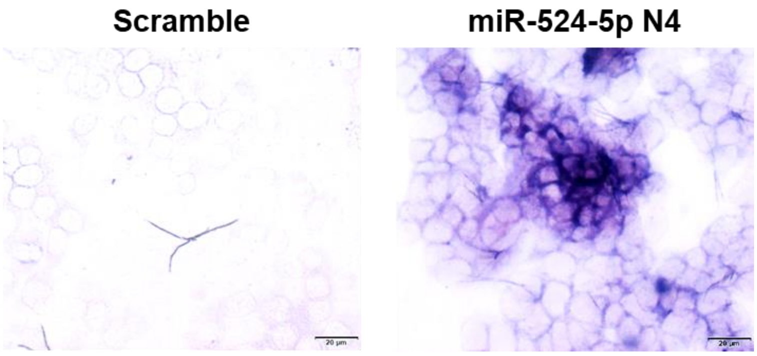


**Fig. S5.** *In situ* hybridization (ISH) of miRNA miR-524-5p in HCT116 cells.

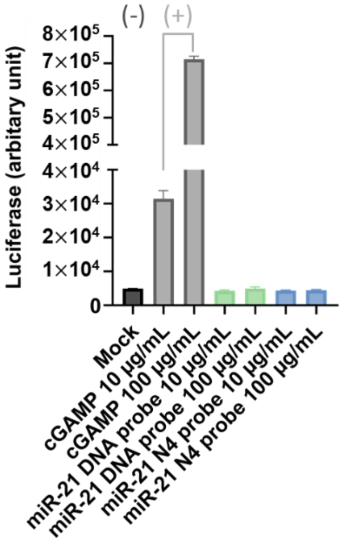


**Fig. S6.** Innate immune response of canonical DNA and antisense N4 nDNA probes. Luciferase activity of the THP1-Dual™ KI-hSTING-R232 reporter cell line treated with the indicated amount of cGAMP, DNA probe or N4 nDNA probe for 24 h.

**Supplemental Tables**

**Table S1.** Intracellular miR-21 concentration

| [a]Human somatic adult cell volume | [b]Number of miRNA copies per somatic adult cell |
|---|---|
| 5,000 $\mu m^3$ | 700–12,000 copies/human genome |
| $5 \times 10^{-15}$ $m^3$ | $1.66 \times 10^{-20}$ mol |
| $5 \times 10^{-12}$ L | $1.66 \times 10^{-11}$ nmol |
| | **Intracellular miR-21 concentration** |
| | ~0.2–4 nM |

a. [2]

b. [3]

**Table S2.** p-value of RT-qPCR with/out DNase I

| | **Unpaired t-test** | |
|---|---|---|
| **Model** | **Heated** | **Unheated** |
| N4 vs DNA | 0.0391 (*) | 0.0001 (***) |
| N4 vs scramble | 0.0018 (**) | <0.0001 (****) |
| N4 vs probe free | 0.0009 (***) | 0.0026 (**) |
| DNA vs scramble | 0.0061 (**) | 0.0046 (**) |
| DNA vs probe free | 0.0018 (**) | 0.0314 (*) |
| scramble vs probe free | 0.0077 (**) | 0.0753 (ns) |

### Appendix

p-value of RT-qPCR with/out DNase I

| Model | Heated vs Unheated |
| --- | --- |
| N4 | 0.7585 (ns) |
| DNA | 0.0489 (*) |
| scramble | 0.4486 (ns) |
| Probe free | 0.8462 (ns) |

p-value of different probe

| Model | Unpaired t-test p-value |
| --- | --- |
| N4 vs DNA | <0.0001 (****) |
| N4 vs LNA | 0.0006 (***) |
| N4 vs scramble | <0.0001 (****) |
| N4 vs probe free | <0.0001 (****) |
| N7 vs DNA | <0.0001 (****) |
| N7 vs scramble | 0.0027 (**) |
| N7 vs probe free | 0.0245 (*) |
| All N vs DNA | 0.0013 (**) |
| All N vs scramble | 0.0016 (**) |
| All N vs probe free | 0.0051 (**) |
| LNA vs DNA | <0.0001 (****) |
| LNA vs scramble | <0.0001 (****) |
| LNA vs probe free | <0.0001 (****) |
| DNA vs scramble | <0.0001 (****) |
| DNA vs probe free | <0.0001 (****) |
